# Proteomics profiling reveals regulation of immune response to *Salmonella* Typhimurium infection in mice

**DOI:** 10.1101/2022.02.08.479500

**Authors:** He Huang, Zachary Even, Alyssa Erickson, Ling Li, Mikhail Golovko, Svetlana Golovko, Ramkumar Mathur, Xusheng Wang

## Abstract

Regulation of the immune response to *Salmonella* typhimurium (*S*. Typhimurium) infection is a complex process, influenced by genetic and environmental factors. Different inbred mouse strains show distinct levels of resistance to *S*. Typhimurium infection, ranging from susceptible (e.g., C57BL/6J) to resistant (e.g., DBA/2J) strains. However, the underlying molecular mechanisms contributing to the host response remain elusive. In this study, we present a comprehensive proteomics profiling of the spleen tissue from C57BL/6J and DBA/2J strains with different doses of *S*. Typhimurium infection by tandem tag mass coupled with two-dimensional liquid chromatography-tandem mass spectrometry. We identified and quantified 3,986 proteins, resulting in 475 differentially expressed proteins between C57BL/6J and DBA/2J strains. Functional enrichment analysis revealed that the mechanisms of innate immune responses to *S*. Typhimurium infection are associated with several signaling pathways, including the interferon signaling pathway. Our proteomic data also discovered a plausible gene in a genomic region that control different levels of resistance to *S*. Typhimurium infection. We further revealed the roles of macrophage cells and pro-inflammatory cytokines in the mechanisms under the natural resistance to *S*. Typhimurium. In summary, our results provide new insights into the genetic regulation of the immune response to *S*. Typhimurium infection in mice.

**Author Summary:** *Salmonella* infection (salmonellosis) is a common zoonotic disease that mainly propagates through contaminated food and drink. Various mouse strains display prominent disparities in responses to *Salmonella* invasion. Elucidating the heterogeneous immune reactions between different mouse strains can shed light on the fundamental molecular mechanisms of the innate immune system. Here, we employed a combination of proteomics and systems biology approaches to provide an unprecedented panorama of the inextricably interlaced immune signaling pathways in response to *Salmonella* infection in mice. Our results revealed the dynamics of cell signaling molecules elicited by inflammation and established new connections among them. We also identified a new candidate gene involved in the combat with pathogens. Our proteomic data and results contribute to understanding the intricate interactions of immune responses to *Salmonella* infection from molecular to systemic levels.

## Introduction

*Salmonella enterica* is a Gram-negative bacterial species that comprises more than 2,600 antigenically different serovars[1]. Some of them, for instance, *Salmonella enterica* serovar Typhimurium (*S*. Typhimurium) can invade a broad range of hosts[2]. Disease manifestations caused by *Salmonella* can range from self-limiting gastroenteritis to a systemic enteric fever[3]. According to the Centers for Disease Control and Prevention (CDC) website[4], *Salmonella* bacteria cause about 1.35 million infections, 26,500 hospitalizations, and 420 deaths in the United States every year. In addition, cross-infection between humans and farmed animals, including chickens, pigs, and cattle, is an important cause of human salmonellosis, especially as the infection can be asymptomatic sometimes[2].

The innate immune to *S*. Typhimurium infection is a complex process and has been proven to be genetically controlled, involving the genomes of the host and pathogen, as well as the environment[5]. *S*. Typhimurium adopts Type III secretion systems (T3SSs) to avoid innate immune receptors and limit the inflammatory response[6]. It has been well known that different inbred mouse strains showed distinct levels of resistance to *S*. Typhimurium infection: susceptible (e.g., C57BL/6J, BALB/cJ, and C3H/HeJ strains), intermediate to resistant (e.g., DBA/2J and A/J strain)[7]. The BXD recombinant inbred (RI) mice from a cross of C57BL/6J (B6) and DBA/2J (D2) inbred strains have been traditionally used to study complex traits under genetic control[8]. Previous studies have shown that the BXD RI mice exhibit substantial variations in the level of virulence to *S*. Typhimurium infection[9]. The two parental strains, B6 and D2, display high susceptibility and very resistance, respectively[7]. However, the underlying molecular mechanism contributing to the immune responses to *S*. Typhimurium infection remains elusive.

Recently developed liquid chromatography coupled with tandem mass spectrometry (LC-MS/MS) has become a powerful technology for large-scale identification of peptides and quantification of protein abundance, allowing us to comprehensively characterize protein expression profile at the genome-wide scale and systematically elucidate molecular mechanisms contributing to the immune responses to *S*. Typhimurium infection. Although RNA-seq profiling of transcriptome in mice after *S*. Typhimurium infection have been conducted[10,11], transcript abundance might not be proportionally correlated to the level of proteins due to various post-transcriptional and post-translational processes[12]. Proteomics is a useful technology complementary to transcriptomics as proteins are the direct coordinators of biological processes. Moreover, coupled with the systems biology approach, proteomics analysis enables generating a panoramic view of the fundamental picture of genetic regulation.

To understand the underlying pathogenic molecular mechanisms of *S*. Typhimurium infection, we perform comprehensive proteomic profiling of the spleen tissue from D2 and B6 strains after *S*. Typhimurium infection by LC-MS/MS, followed by integrative analysis to reveal differential expression proteins between B6 and D2 strains and signaling pathways related to the immune response to *S*. Typhimurium infection. We next discover a new candidate gene for a previous unsolved quantitative trait locus (QTL) that is responsible for the divergent immune responses. We also conducted a macrophage phagocytosis assay and quantification of pro-inflammatory cytokines.

## Results

### Proteome profiling of mouse spleen after *S*. Typhimurium infection

To profile and systematically characterize the changes of protein expression after *S*. Typhimurium infection in mice, we used multiplexed Tandem Mass Tag (TMT) and LC/LC-MS/MS strategy to quantify the proteome of spleen from B6 and D2 mice 14 days after *S*. Typhimurium infection with high dosage (5×10^8^ CFU) and low dosage (5×10^6^ CFU). Spleen tissue samples were lysed, digested, and labeled with different TMT tags, then pooled and analyzed by LC/LC-MS/MS (**Fig 1A**). A total of 3,986 proteins derived from 20,270 peptides were identified and quantified at the protein FDR of 1% (**S1 Table**). Principal-component analysis (PCA) showed high reproducibility between two biological replicates (i.e., male and female), but a clear separation between control and infection groups and between B6 and D2 strains (**Fig 1B**). To differentiate experimental from biological variances, we compared intraclass (i.e., replicates) and interclass (i.e., strains) correlations (**Fig 1C**). The two biological replicates in the intraclass group displayed a high correlation, whereas two different mouse strains exhibited considerable variation in protein expression after *S*. Typhimurium infection.

**Fig 1.**
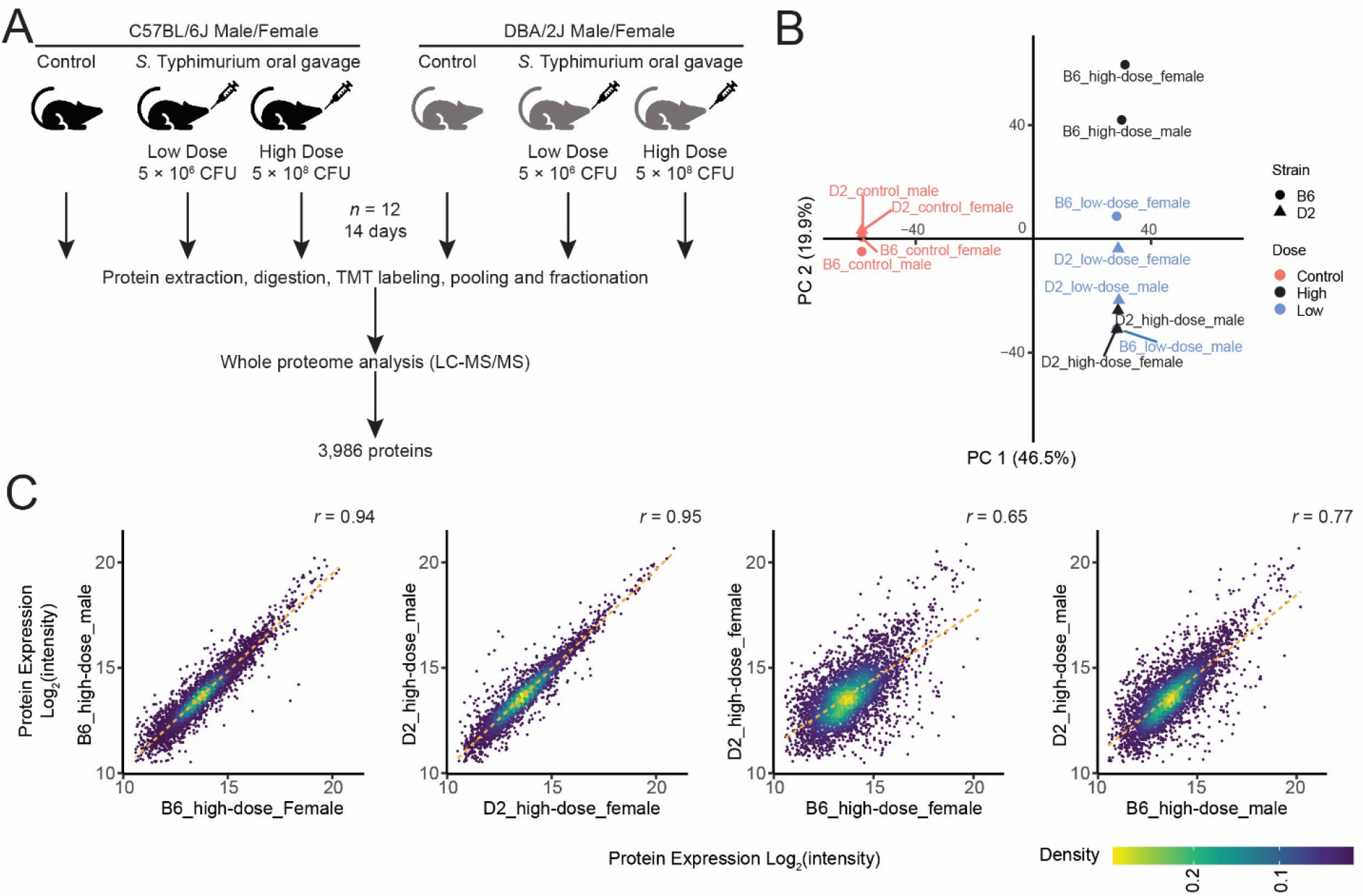
Proteomics profiling of C57BL/6J (B6) and DBA/2J (D2) mouse spleen tissues from *Salmonella enterica* serovar Typhimurium (*S*. Typhimurium) infection and control groups. **A.** Experimental design: B6 and D2 mice were infected with *S*. Typhimurium strain 14028GFP by oral administration of *S*. Typhimurium at two dosages (5×10^6^ and 5×10^8^ CFU, colony-forming unit) for 14 days for male and female individual of each mouse strain. Whole proteome analysis was performed on liquid chromatography-tandem mass spectrometry (LC-MS/MS). **B.** Principal component analysis showed high correlation between biological replicated and clear separation between control and infection groups. **C.** Comparison of interclass and intraclass correlations differentiated experimental from biological variances.

The most prominent regulator of the susceptibility to *Salmonella* was identified as *Ity* (immunity to Typhimurium), a dominant autosomal gene located at chromosome 1 [13–15]. This *Ity* controlled innate resistance immediately prior to the exponential growth phase was mediated by a localized response in the spleen, more specifically the macrophages [16,17]. Consistent with the previous observation, we observed that ITY protein expression in D2 strains is significantly elevated compared to that in B6 strains after both high dose and low dose infection (**S1 Fig**).

### Differentially expressed proteins between B6 and D2 in the mouse spleen after *S*. Typhimurium infection

To detect differentially expressed proteins between B6 and D2 strains, we performed differential protein expression analysis using a linear model with the limma package[18]. We detected 118 differentially expressed proteins (DEPs) from the interaction effects of strain and dosage less than 5% FDR (**Fig 2A**, **S2 Table**). Using the limma package, in high dose group, we identified 475 DEPs between B6 and D2 mice (5% FDR, fold change cut-off of 1.5; **Fig 2B**, **S3A Table**). Enrichment analysis showed that these DEPs between strains were involved in various small molecule catabolic and metabolic processes (**Fig 2C**). We next sought to characterize alterations in protein expression in response to different dosages of infection. In B6 mice, we identified 189 DEPs between high dose and control group, and 178 DEPs between high dose and low dose group (**Fig S2**, **S3B-C Table**). In contrast, in D2 mice, no DEPs were detected among control, low dose, and high dose groups. In the following analysis, we focused on the 475 DEPs between B6 and D2 strains for subsequent analyses.

**Fig 2.**
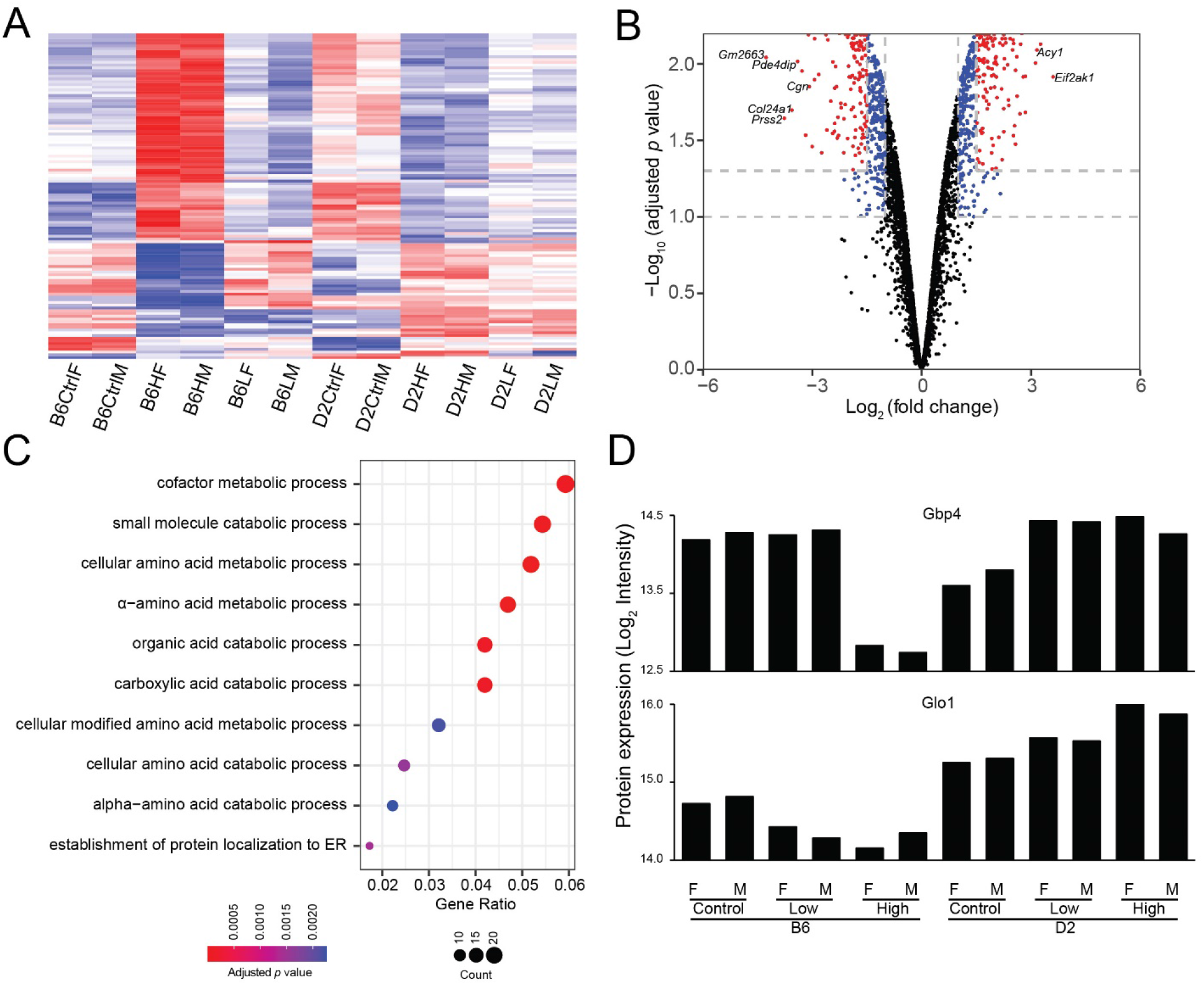
Differential protein expression analysis. **A.** 118 differentially expressed proteins (DEPs) were detected from the interaction effects of strain and dosage by two-way ANOVA (< 5% FDR). **B.** In high dose (high dose) infection group, we identified 475 DEPs between B6 and D2 mice (5% FDR, fold change cut-off of 1.5). **C.** Enrichment analysis showed that the DEPs between strains in high dose group were involved in various small molecule catabolic and metabolic processes. **D.** Two representative DEPs, GBP4 (Guanylate-binding protein 4) and Glo1 (glyoxalase I) displayed distinctive expression patterns among infection and control groups from B6 and D2 mouse strains.

A notable example is guanylate-binding protein 4 (GBP4) (**Fig 2D**), an IFN-γ-inducible GTPase protein [19], which is indispensable for inflammasome activation and *S.* Typhimurium clearance. Our quantification data showed that high-dose *S*. Typhimurium infection suppressed GBP4 expression significantly in B6 strain although low-dos infection did not change GBP4 expression substantially. In contrast, the expression level of GBP4 was elevated in both low- and high-dose *S*. Typhimurium infection groups in D2 mice. This contrasting pattern in *Salmonella* susceptible and resistant strains implies that GBP4 might be an imperative factor to which the differences in natural resistance to *Salmonella* might be attributed. A previous study in Zebrafish reported GBP4-dependent resistance to *S*. Typhimurium mediated by the prostaglandin D2 pathway of neutrophils[20]. However, seemingly contradicting evidence pointed to the suppression effects of innate immune response to viruses by interrupting the interaction of TNF receptor-associated factor (TRAF) family members TRAF6 with Interferon Regulatory Factor 7 (IRF7), causing impairment of Type I IFN (IFN-α) expression[21].

Another example is glyoxalase I (GLO1), a ubiquitously expressed enzyme involved in the methylglyoxal detoxification[22]. Methylglyoxal detoxification metabolism is required for the survival of *S*. Typhimurium in the mammalian host[23]. Our result revealed that the expression of GLO1 was significantly lower in B6 mice than that in D2 mice (**Fig 2D**). Moreover, the expression patterns of GLO1 expression among control, low dose, and high dose in B6 and D2 strains changed in opposite directions, with down-regulation in B6 but upregulation in D2.

### Co-expression network analysis unveils genetic control of mouse innate resistance to *Salmonella*

To reveal genetic control of innate resistance to *S*. Typhimurium infection, we identified co-expression clusters of functionally related proteins by weighted gene correlation network analysis (WGCNA) (**Fig 3A**). Three co-expression protein clusters were identified from 475 DEPs that showed the difference in protein expression between B6 and D2 strains (**Fig 3B and 3C, S4 and S5 Tables**). Cluster 1 contains 214 (45%) DEPs whose expression displayed upregulation in B6 compared to D2 with high-dose infection. In contrast, the expression of these proteins remained at similar levels in B6 and D2 with low-dose infection. Enrichment analysis showed that most of these were involved in the turnovers of amino acids and proteins, including the metabolic and catabolic processes of cellular amino acids and post-translational transportation of proteins (**Fig 3C**), indicating the possible mobilization of protein synthesis machinery in response to invasion of pathogen.

**Fig 3.**
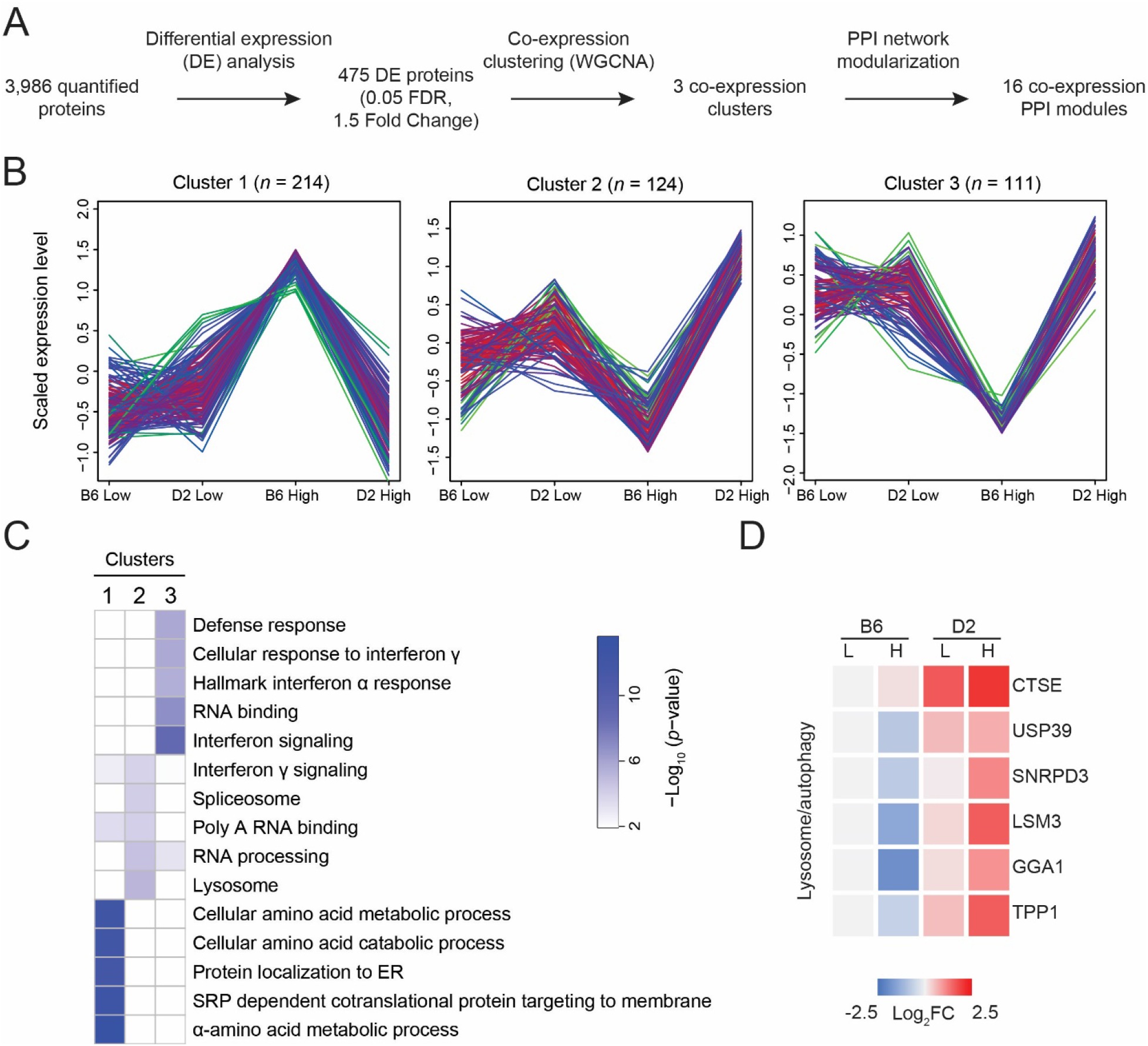
Co-expression analysis. **A.** Weighted gene correlation network analysis (WGCNA) followed by protein-protein interaction (PPI) network modularization. **B.** Three co-expression protein clusters were detected from 475 DEPs that showed strain difference between B6 and D2 after high dose infection. **C.** The pathways involved in the three co-expression clusters. **D.** Cluster 2 contains a group of lysosome and autophagy pathway-related proteins that were significantly upregulated in D2 mice compared to those in B6 ones.

Conversely, cluster 2 showed noticeable downregulation in B6 compared to D2 after high-dose infection. Cluster 2 is composed of 124 (26%) DEPs, which are primarily responsible for RNA binding and processing. In comparison to the *Salmonella* susceptible B6 strain, the proteins in cluster 2 slightly upregulated after low-dose infection and pronounced increase in the *Salmonella* resistant D2 strain. The patterns of change in protein expression in cluster 1 and cluster 2 suggested that the innate immune response of D2 strain to *Salmonella* infection might be predominantly taking place at the level of RNA regulation. In B6 mouse, the unleashed rapid proliferation of *Salmonella* in B6 mice might deploy large amount of protein synthesis machinery.

Further scrutinizing DEPs in cluster 2, we observed a group of lysosome and autophagy pathway-related genes that were significantly upregulated in D2 mice compared to those from B6 mice (**Fig 3D**). It has been well established that autophagosomes selectively targeted *S*. Typhimurium and delivered the pathogens to lysosome for degradation as an antibacterial defensive mechanism[24]. Our results indicated that the activation of lysosome/autophagy pathway may account for the resistance to *Salmonella* for D2 mice. Noticeably, our results demonstrated the expression of CTSE gene, encoding an aspartic proteinase (Cathepsin E) necessary for class II antigen presentation in macrophages[25], markedly raised in D2 after *Salmonella* infection compared to B6.

Cluster 3 comprised 111 (23%) of the DE proteins appertaining to the pathway of immune response. Noticeably, many major histocompatibility complex (MHC) class II molecules were identified among these DEPs in this cluster. MHC class II molecules have been well known for presenting antigens derived from exogenous sources, including bacteria to CD4+ T cells, in contrast to MHC class I molecules, presenting peptides originated from intracellular sources to CD8+ cells[26]. The pathway analysis implied that many DEPs in cluster 3 function in the interferon (IFN) signaling pathway.

### Protein-protein interaction network analysis suggests signaling pathways involved in innate immune response

To further identify signaling pathways involved in innate immune response to *S*. Typhimurium infection, we superimposed proteins within each cluster onto the PPI network, identifying 16 functional modules (**Fig 3A**; **S6 Table**). Each module consisted of 2 to 28 proteins (**S6 Table**). Many of these modules were associated with known signaling pathways (**S6 Table**). Prominently, module 2 in cluster 3 contained 13 genes of IFN signaling pathway (**S6 Table**; **Fig 4A**), including GBP1, GBP2, GBP4, GBP5, ISG20 (Interferon Stimulated Exonuclease Gene 20), IRF3 (Interferon Regulatory Factor 3), STAT1 (signal transducer and activator of transcription 1), PARP9 (Poly (ADP-Ribose) Polymerase Family Member 9) (**Fig 4B**). Previous studies showed that loss-of-function mutations in the IFN signaling pathway resulted in vulnerability to infections by various pathogens, including *Salmonella*[19]. The expressions of hundreds of genes were elicited by IFN signaling pathway, inclusive of p47 and p65 GBPs[19]. A recent research unveiled that after Gram-negative bacteria (*Salmonella, Shigella*) infection, interferon-induced GBPs 1-4 were assembled on the surface of bacteria in a hierarchical manner, which further activate caspase-4 and its downstream pyroptotic program mediated by Gasdermin-D (GSDMD) and IL-18[19].

**Fig 4.**
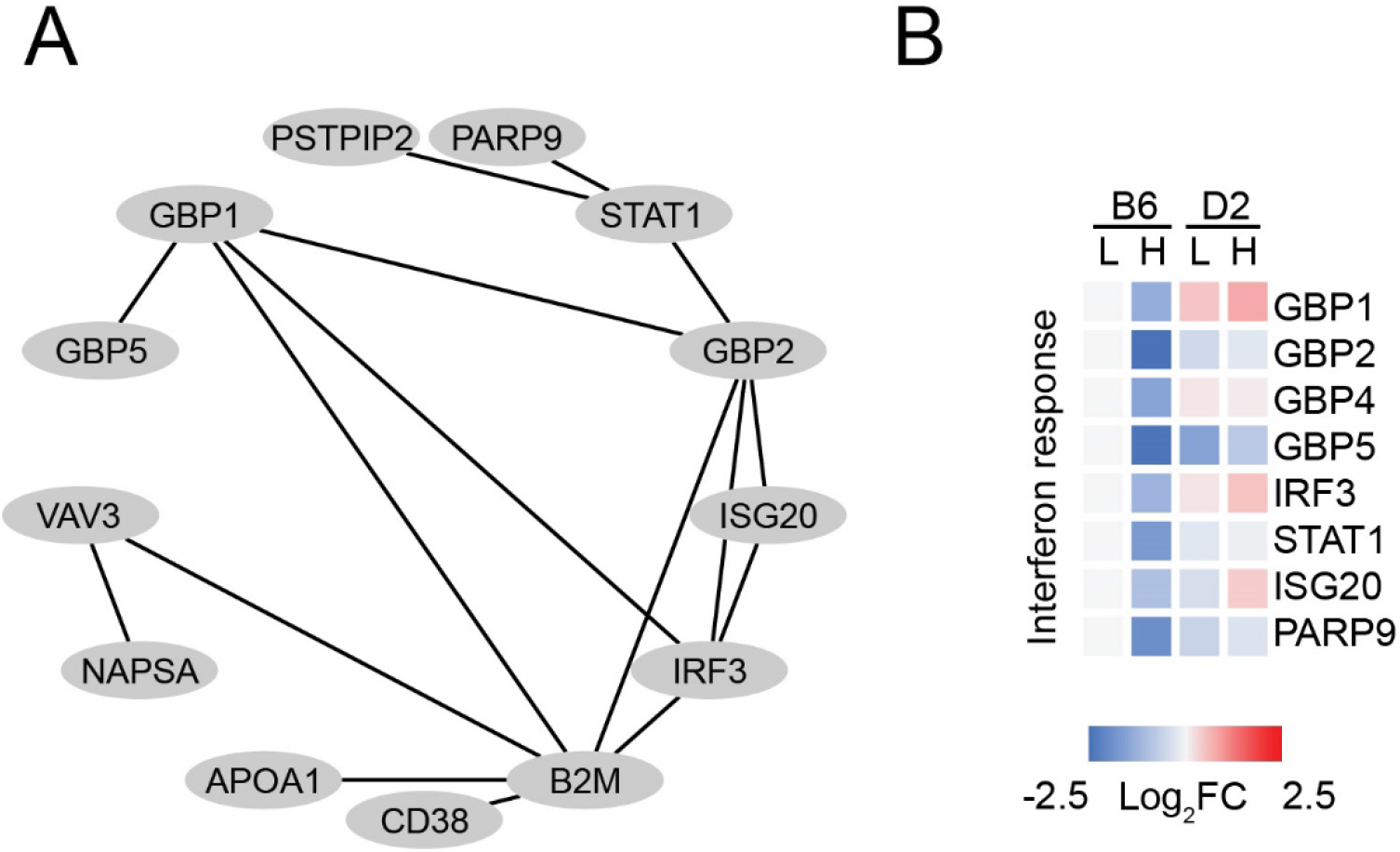
Functional modules were identified by superimposition of co-expression clusters on the PPI network. **A.** The interaction network build on the 13 genes of IFN signaling pathway in module 2 of cluster 3. **B.** Proteins involved in IFN signaling pathway showed different pattern of expression in B6 and D2 mice between infection and control groups.

Another notable example is module 1 in cluster 1, the largest module consisting of 28 DEPs. Most of these DEPs were involved in the protein synthesis and post-translational pathway, accounting for the changes at translational level after *S*. Typhimurium infection. Correspondingly, module 1 in cluster 2, the second largest module, was mainly composed of DEPs enriched in RNA synthesis, splicing, processing, and binding, which reflected the responses of organisms to *S*. Typhimurium infection at transcriptional level. Altogether, our results of the network analysis provided a panoramic picture of the coordinated immune and inflammatory mechanisms after *S*. Typhimurium infection in mice.

### Proteomics data reveals a candidate gene under a QTL interval

QTL mapping in BXD mice has previously identified that 72.4 – 73.4 Mb of chromosome 1 is a genomic locus responsible for different levels of resistance to *S.* Typhimurium infection[9] (**Fig 5A and 5B**). However, the candidate genes have not yet been identified. We postulate that integration of our proteomic data allows us to facilitate the identification of candidate genes within the QTL interval. Our proteomic profiling data detected several genes in this QTL interval, including *Smarcal1, Rpl37a, Tnp1*, and *Pinc* (**Fig 5B**). The expression of RPL37A (60S ribosomal protein L37a) protein showed a significant difference between B6 and D2 after high-dose infection (**Fig 5C**). Intriguingly, although RPL37A was typically considered as a classic housekeeping gene for normalizing protein expression, studies have shown that ribosomal proteins play crucial roles in immune response to pathogen infections[27,28]. Our results indicated that RPL37A could be a plausible candidate gene for this QTL.

**Fig 5.**
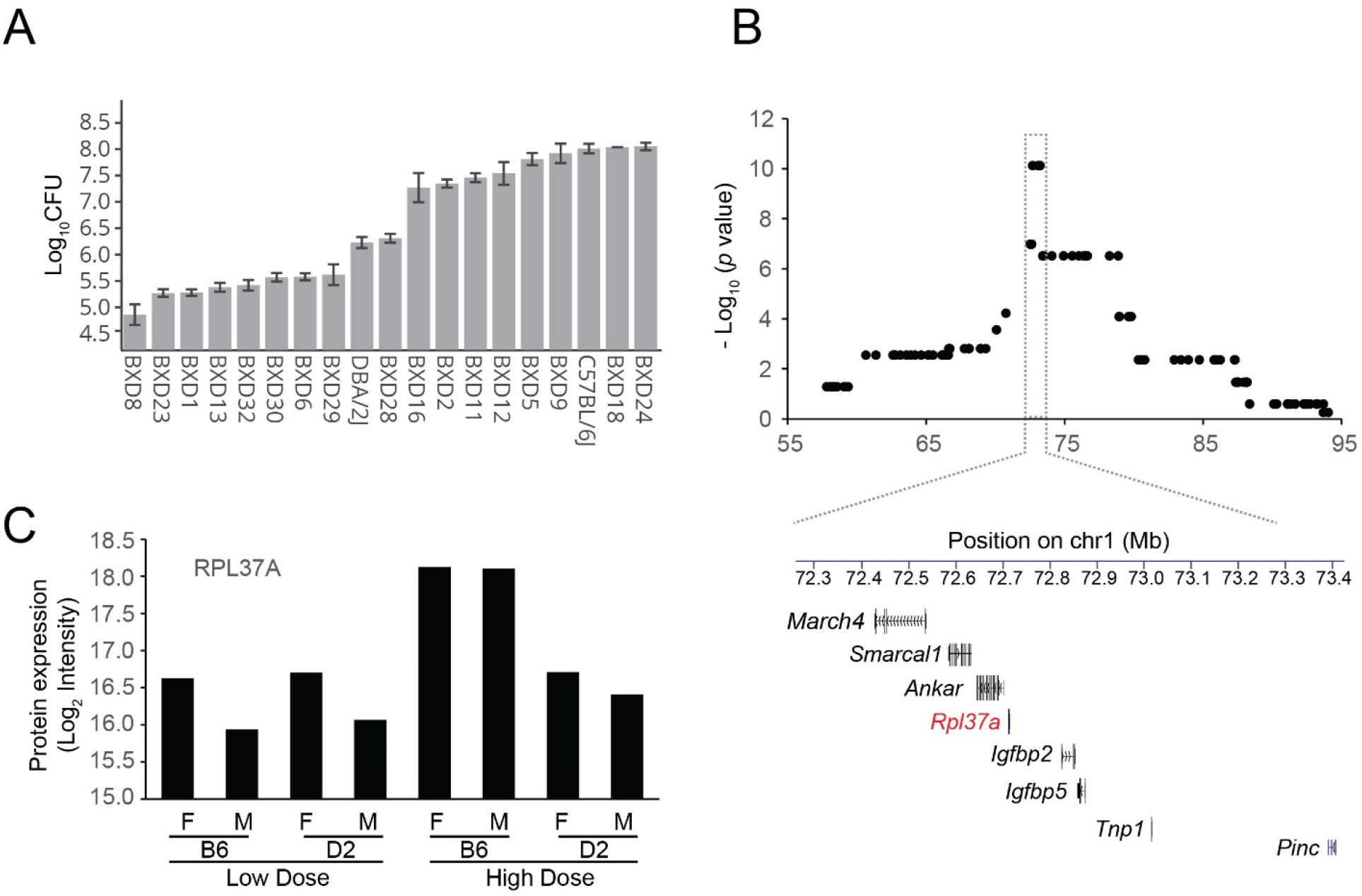
Proteomics data reveals candidate genes under QTL interval. **A.** Quantitative trait locus (QTL) mapping of the different levels of resistance to *S*. Typhimurium infection in BXD mice[9]. **B.** Our proteomic profiling data detected several genes in the QTL interval that determined the different levels of resistance to *S*. Typhimurium infection (72.4 - 73.4 Mb of chromosome 1). **C.** RPL37A (60S ribosomal protein L37a) protein showed a significant disparity between B6 and D2 mice after high dose of *S*. Typhimurium infection.

### Macrophage cells and pro-inflammatory cytokines in host defense against *S*. Typhimurium infection

To understand how macrophage is involved in *S*. Typhimurium infection, we performed macrophage phagocytosis assay and experimental validation of cytokines in B6 and D2. Our results demonstrated that the macrophage cells from D2 mice were able to clear *Salmonella* and better than those from B6 mice (**Fig 6A)**. Many intracellular bacteria, represented by GFP fluorescent signaling, were found in the macrophages from B6 whereas very few existed in those from D2. This disparity indicates the attribution of the ability of macrophage to kill invaded pathogens to the innate immune responses to *S*. Typhimurium infection. Our RT-qPCR quantification of the cytokines in the macrophages from two mouse strains further revealed strikingly different pattern of expression (**Fig 6B**). Some pro-inflammatory cytokines, including IFN-γ and iNOS (inducible nitric oxide synthase), were significantly upregulated in D2 mice, but not in B6. In contrast, other cytokines (e.g., IFN-β, IL-6, and IL-8) were significantly upregulated in B6 mice, but not in D2.

**Fig 6.**
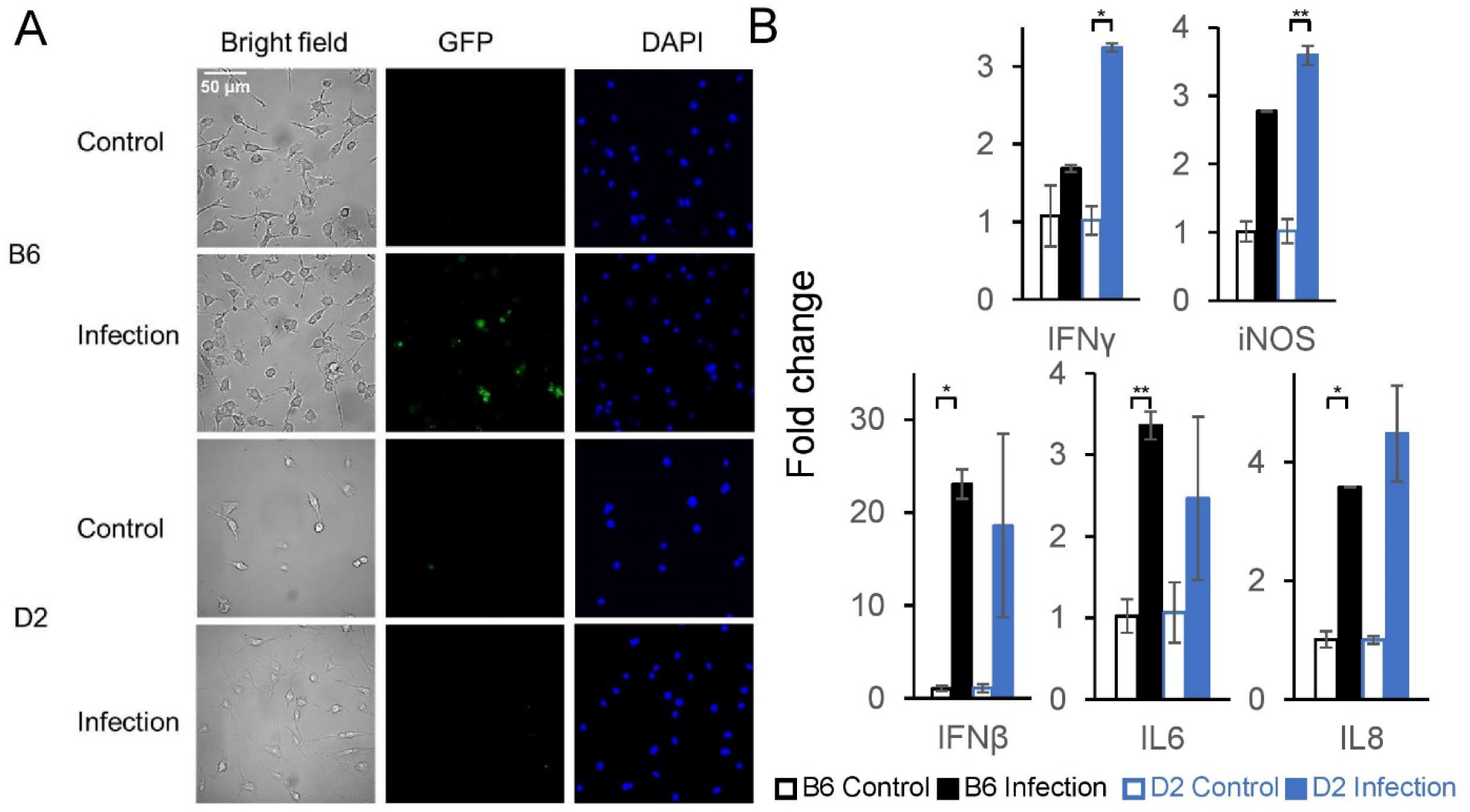
Macrophage cells and pro-inflammatory cytokines play important parts in host defense against *S*. Typhimurium infection. **A.** Macrophage phagocytosis assay revealed better clearance of pathogens by macrophages from D2 mice than those from B6 ones. **B.** Different sets of pro-inflammatory cytokines were elicited in B6 and D6 mice after *S*. Typhimurium infection.

## Discussion

In this study, we conducted proteomic profiling to understand the mechanisms responsible for the natural resistance to *Salmonella*. Our analysis identified ~4,000 proteins, 475 of which displayed significantly different levels of protein expression between B6 and D2 mice after *S*. Typhimurium infection. By combining our proteomics data and linkage analysis, we identified RPL37A as a plausible candidate gene for a QTL that influences the host response to *S*. Typhimurium infection in mice. We experimentally validated macrophage cells and pro-inflammatory cytokines that are involved in host defense against *S.* Typhimurium infection. Our results presented here provide an integrative network accountable for natural resistance to *Salmonella*, connecting the functionalities of individual genes and pathways for a panoramic view of the molecular mechanism underlying *S.* Typhimurium infection.

Our study, for the first time to the best of our knowledge, applied a large-scale proteomic strategy to reveal the genetic controls of innate immune response to *Salmonella* by comparing the differential protein expression profiles of *Salmonella* resistant and susceptible mouse strains. We identified 475 DEPs and several signaling pathways involved in host defense against salmonella infections. Our network analysis strongly pointed to the key functions of IFN signaling pathway in the natural resistance to *S*. Typhimurium infection by identifying multiple protein elements, including GBPs, STAT1, IRF3, ISG20 (**Fig 4**). Although IFNs were typically considered as pleiotropic cytokines in anti-viral gene program, plenty of studies have indicated its role in host defense against bacterial infections[29]. Our results strongly exhibited the upregulation of IFN-γ (type II IFN or immune IFN) related proteins in *Salmonella* resistant D2 strain compared to B6 strain. It has been well established that IFN-γ worked as a protective cytokine in the natural host response to several bacterial infections[30]. The important role of IFN-γ in innate host response was supported by the significantly compromised immune responses in mutant mice with gene disruptions in IFN-γ signaling pathway[30]. In addition, MHC molecules, one of the most predominant upregulated proteins in response to *Salmonella* infection in cultured macrophages and intestinal epithelial cells, were regulated by IFN-γ as well[30]. On the other hand, interestingly, there was evidence of *S*. Typhimurium induced production of type I interferon leading to necroptosis of macrophages, allowing the pathogens to escape the host immune response[31]. Further study demonstrated that host type I IFN system facilitated the spread of *S*. Typhimurium by IFN-β–mediated repression of macrophage innate immune transcriptional responses[32]. We speculated that to counteract the natural resistance to *Salmonella* in D2 strain, the pathogens might strategically mobilize more proteins involved in the type I IFN system.

Both IFNα (type I IFNs) and IFN-γ (type II IFNs) activates the classical JAK (Janus activated kinase)-STAT (signal transducer and activator of transcription) pathway of signaling[33]. Our results suggested the upregulation of type I IFNs in B6 mice whereas type II IFNs in D6 mice. This difference may account for the distinctive innate immune response to *Salmonella* infection. Since the activation of JAK-STAT signaling pathway involves the changes of phosphorylation status of its components, further phosphoproteomics study may reveal the detailed regulatory mechanisms. iNOS plays an important part in the polarization of macrophages between M1 (pro-inflammatory) and M2 (anti-inflammatory)[34]. IFN-γ and iNOS can differentiate macrophages into M2 macrophages and therefore promote inflammation and the expression of iNOS depends on IFN-γ[35]. The upregulation of the pro-inflammatory cytokines such as, IFN-γ and iNOS in the macrophages from D2 mice implied the importance of inflammation process (the polarization to M1 macrophages) in the innate immune response to *Salmonella* infection.

Our results not only integrated many previously studied proteins in the macrophage-mediated immune response, including components in the IFN signaling pathway, but also suggested novel functions of some proteins. For example, our data demonstrated a strikingly different expression pattern of Glo1 between D2 and B6 mice (**Fig 2D**). Glo1 was found to be responsible for converting cytotoxic metabolite methylglyoxal (MG) to S-D-lactoylglutathione, previously identified as a crucial component in osteoclastogenesis[22]. The receptor activator of NF-κB ligand (RANKL) and macrophage colony stimulating factor (M-CSF) were recognized as key regulators for osteoblasts in this process. The expression of Glo1 was detected in the macrophages in mice[36]. Interestingly, a recent study has indicated the role of Glo1 in alleviating the immune-inflammatory response by regulating the apoptosis pathway mediated by TNF-α[37]. Consider the importance of RANKL and M-CSF in the immune response; it would be of great interest to further decipher their enigmatic roles in the natural resistance to *S*. Typhimurium infection.

Contradictory to our result showing the increased expression of RPL37A protein in B6 mice after high dose *S*. Typhimurium infection (**Fig 5C**), a previous study demonstrated that in porcine neutrophil, RPL37 was downregulated after *Salmonella* infection, which resulted in the inhibition of TNF pathway and therefore promoted *Salmonella* survival and replication[27]. We suspect that in B6 mice, the rising of RPL37A expression may server as a complementary mechanism in response to the relatively ineffective immune reaction to *Salmonella* invasion. Another study suggested the important functions of 60S ribosomal proteins in the presentation of MHC Class I antigenic peptides using recombinant influenza A virus construction[28]. Further inquiries into the potential roles of RPLs in the generation of MHC class II peptides would be a riveting direction.

Although we identified proteins of a series of related signaling pathways and cellular machinery, our current analysis has several limitations. First, the analysis was limited by the depth of proteomic data obtained, which can be improved by employing more fractionations. Second, our proteomic measurement was based on a bulk tissue, which only reflects the mean expression of different cell populations. Single-cell proteomics technologies, such as SCoPE-MS (Single Cell ProtEomics by Mass Spectrometry) and nanoPOTs (Nanodroplet Processing in One Pot for Trace Samples), can be used for measuring cell-type-specific expression. This would be especially useful in characterizing the roles of different immune cells in the host defense response. Third, integrating proteomic and transcriptomic data will further improve our understanding of the genetic basis of the host response to *Salmonella* infection.

In summary, our proteomics profiling analysis together with quantification of pro-inflammatory cytokines and phagocytosis assay unveiled the connect-the-dots explanation for the different immune responses between D2 and B6 mouse strains, integrating multiple cell signaling pathways. We also identified novel roles of several proteins in the innate immune system. Our proteomic data and systems biology approaches provide an opportunity to understand the intricate processes of immune responses to *Salmonella* infection in mice from molecular to systemic levels.

## Materials and Methods

### Ethics statement

All procedures for mice infection studies were approved by the Institutional Animal Care and Use Committee at the University of North Dakota (IACUC Approval No. 2008-3). Mice were euthanized by carbon dioxide inhalation followed by cervical dislocation.

### Animals

The following mouse strains were used in this study: C57BL/6J (B6) and DBA/2J (D2). Both male and female mice of each strain were used as biological replicates in this study. All animals were purchased from The Jackson Laboratory. Animals were housed and maintained in a 12:12 light/dark cycle, with *ad libitum* access to food and water. Mice at 12-week-old were infected with *S*. Typhimurium strain 14028GFP by oral administration of *S*. Typhimurium at two dosages (5×10^6^ and 5×10^8^ CFU, colony-forming unit) for 14 days for male and female individuals of each mouse strain. A total of 12 mice were used for the experiment. *S. Typhimurium* was grown in Luria Bertani (LB) medium at 37 °C. The bacterial titer was determined by a standard CFU count on LB agar plate. Mice will be orally inoculated with *S. Typhimurium* by gavage needle. Mice were assayed daily for morbidity (determined as % weight loss), mortality, and clinical disease scores. After 14 days post-infection, mice were sacrificed and dissected rapidly. Spleen samples were collected, frozen in liquid nitrogen, and stored at −80°C for the subsequent proteome profiling.

### Protein extraction and quantification

The frozen spleen samples were weighed and homogenized in the lysis buffer (50 mM HEPES, pH 8.5, 8 M urea, and 0.5% sodium deoxycholate, 100 μL buffer per 10 mg tissue). Samples were lysed in Bullet Blender (Next Advance) under speed 8 for 30 s at 4°C, rested for 5 s, and repeated 5 times or until the samples were homogenized. Protein concentration was measured by the BCA assay (Thermo Fisher).

### Protein digestion and tandem-mass-tag (TMT) labeling

Quantified protein samples (200 μg in the lysis buffer with 8 M urea) for each TMT channel were treated by 1 mM dithiothreitol (DTT) for 1 hour at room temperature to reduce disulfide bonds. Samples were diluted to 2 M urea with 50 mM HEPES (pH 8.5) and proteolyzed with trypsin (Promega, 1:50 w/w) at room temperature overnight. Peptides were reduced by 1 mM DTT for 2 hours at room temperature and then incubated with 10 mM incuiodoacetamide (IAA) for 30 minutes in the dark to alkylate Cys-containing peptides. The reaction was quenched by 30 mM DTT for 30 minutes at room temperature. Samples were acidified by adding trifluoroacetic acid (TFA) to achieve 1% concentration and then centrifuged at 20,000 × g at room temperature for 10 minutes. The supernatant was collected and desalted by the Sep-Pak C18 cartridge (Waters), and then dried by centrifugal vacuum concentrator. Each sample was resuspended in 50 mM HEPES (pH 8.5) for TMT labeling following the manufacturer’s protocol and then mixed equally, followed by desalting for the subsequent fractionation.

### Extensive two-dimensional liquid chromatography-tandem mass spectrometry (LC/LC-MS/MS)

The TMT labeled samples were fractionated by offline basic pH reverse phase LC, yielding 20 fractions. Each fraction was analyzed by the acidic pH reverse phase LC-MS/MS. In the acidic pH LC-MS/MS analysis, each fraction was run sequentially on a column (75 μm ID x 20 cm L x 365 μm OD, CoAnn Technologies, LLC, Cat. No. HEB07502001718I) interfaced with a Q Exactive Orbitrap (Thermo Fisher). Peptides were eluted by a four hours gradient (buffer A: water; buffer B: acetonitrile). MS settings included the MS1 scan (445 - 1600 m/z, 70,000 resolution, 1 × 10^6^ AGC and 60 ms maximal ion time) and 10 data-dependent MS2 scans (fixed first mass of 120 m/z, 70,000 resolution, 1 × 10^5^ AGC, 105 ms maximal ion time, 1.0 m/z isolation window).

### Identification of proteins by database search with JUMP software

We used JUMP search engine to search MS/MS raw data against a composite target/decoy database to evaluate FDR. All original target protein sequences were reversed to generate a decoy database that was concatenated to the target database. FDR in the target database was estimated by the number of decoy matches (*nd*) and the number of target matches (*nt*), according to the equation (FDR = *nd*/*nt*), assuming mismatches in the target database were the same as in the decoy database. The target database was downloaded from the UniProt mouse database (59,423 entries), and decoy database was generated by reversing targeted protein sequences. Major parameters included precursor and product ion mass tolerance (±15 ppm), full trypticity, static mass shift for the TMT tags (+304.2071453) and carbamidomethyl modification of 57.02146 on cysteine, dynamic mass shift for Met oxidation (+15.99491), maximal missed cleavage (n = 2), and maximal modification sites (n = 3). Putative PSMs were filtered by mass accuracy and then grouped by precursor ion charge state and filtered by JUMP-based matching scores (Jscore and ΔJn) to reduce FDR below 1% for proteins during the whole proteome analysis. If one peptide could be generated from multiple homologous proteins, based on the rule of parsimony, the peptide was assigned to the canonical protein form in the manually curated Swiss-Prot database. If no canonical form was defined, the peptide was assigned to the protein with the highest PSM number.

### TMT-based peptide/protein quantification by JUMP software suite

Protein expression was quantified using the following steps with JUMP software suite: (i) TMT reporter ion intensities were extracted for each PSM; (ii) the raw intensities were corrected based on isotopic distribution of each labeling reagent; (iii) PSMs with very low intensities (e.g. minimum intensity of 1,000 and median intensity of 5,000) were excluded from the subsequent analysis; (iv) Sample loading bias was normalized with the trimmed median intensity of all PSMs; (v) the mean-centered intensities across samples was calculated, (vi) protein relative intensities by averaging related PSMs was calculated; (vii) protein absolute intensities by multiplying the relative intensities by the grand-mean of three most highly abundant PSMs was computed.

### Principal component analysis

Principal component analysis (PCA) was used to visualize the differences among samples. All gene and metabolite abundance were used as features of PCA. The pairwise Euclidean distance between features was calculated. PCA was performed using the R package prcomp (version 3.4.0).

### Differential expression analysis

Differentially expressed proteins between the two strains were identified by Analysis of Variance (ANOVA) and the limma R package (version 3.46.0). The Benjamini-Hochberg method for false discovery rate correction was used, and proteins with an adjusted p-value < 0.05 and log_2_ (fold change) > 1.0 were defined as differentially expressed.

### Pathway enrichment

To assess the functional relevance of the differentially expressed proteins, the R package clusterProfiler (version 3.18.1) was used for gene ontology enrichment analysis[38]. Gene ontology terms with a Benjamini-Hochberg adjusted *p*-value < 0.05 were defined as significantly enriched.

### Weighted gene correlation network analysis (WGCNA)

WGCNA was performed as described before[39] with the WGCNA R package[40], using an integrated software suite JUMPn[41]. The DE proteins were used to construct the cluster. Pearson correlation matrix (with direction, i.e., for building signed correlation network) was calculated. An adjacency matrix was calculated by raising the correlation matrix to a power of 16 using modified scale-free topology criterion[42]. Correlation clusters were defined by hybrid dynamic tree-cutting method[40] with the minimum height for merging modules of 0.2. A consensus trend was calculated based on the first principal component (i.e., eigengene) for every cluster. Proteins were assigned to the most correlated cluster with a minimum cutoff of Pearson correlation coefficient (*r*) of 0.7.

### Protein-protein interaction (PPI) network analysis

PPI network analysis was conducted as described before[39], using an integrated software suite JUMPn[41]. Proteins in each cluster by WGCNA were superimposed onto a heterogeneous composite PPI database that combined STRING (v10)[43], BioPlex[44], and InWeb_IM[45], which included 18,515 proteins and 469,993 PPI connections. Modules in each protein cluster were defined by a two-step procedure: (i) PPI edges were retained only if both nodes (i.e., the two connected proteins) came from proteins in the same cluster; (ii) a topologically overlapping matrix[46] for the PPI network was computed; and this network was modularized into individual modules by the hybrid dynamic tree-cutting method[40]. Three pathway databases, including Gene Ontology (GO), KEGG and Hallmark by Fisher’s exact test were used to annotate the modules, which were visualized by Cytoscape[47].

### Macrophage phagocytosis assay

B6 and D2 mice, one male and one female individual for each strain, were sacrificed by carbon dioxide (CO_2_) inhalation followed by cervical dislocation. The abdomen regions and hind limbs were sterilized with 70% ethanol. The bone marrow of femur and tibia was harvested by flushing the bones with Dulbecco’s Modified Eagle Medium (DMEM) (Gibco™, Thermo Fisher) using a 1 mL insulin syringe in a laminar flow hood. The bone marrow cells were pipetted up and down to bring the cells into single-cell suspension. Macrophages were differentiated in 10 cm cell culture petri dishes in DMEM supplemented with 10% heat-inactivated fetal bovine serum (FBS) (Gibco™, Thermo Fisher), 30% L929-conditioned medium, and 10% penicillin-streptomycin (10,000 U/mL) (Gibco™, Thermo Fisher) in a humidified incubator with 5% CO_2_ at 37°C. Fresh medium was added every 2-3 days. Macrophage progenitors adhered to the cell dish and were not washed away. Macrophages were fully differentiated by day 6. On the 7^th^ day, macrophage cells were counted using a hemocytometer. One million macrophages were seeded into one well of a 12-well plate, with one coverslip in each well. *S*. Typhimurium bacteria were introduced into the medium with a nominal multiplicity of infection (MOI) of 10. Infection proceeded for 15 minutes. Then, macrophage cells were washed twice with PBS supplemented with 20% penicillin-streptomycin to remove extracellular bacteria and then incubated in DMEM medium for 2 hours. The coverslips were taken out and mounted with Fluoromount-G™ Mounting Medium, with DAPI (4’,6-diamidino-2-phenylindole) (Invitrogen™, Thermo Fisher) on microscope slides. Fluorescent images were obtained by Leica DMi8 Thunder fluorescent microscope (Leica Camera, Wetzlar, Germany), by 550 nm excitation for green fluorescent protein (GFP) expressed in *S.* Typhimurium strain 14028GFP, and 395 nm excitation for DAPI. The rest of the macrophage cells were collected for quantitative reverse transcription PCR (RT-qPCR) analysis.

### RT-qPCR quantification of cytokines

Total RNA was isolated from macrophages using the TRIzol™ Reagent (Thermo Fisher). To extract RNA, macrophage cells were washed with PBS and quickly homogenized in TRIzol™ Reagent. The RNA pellets were resuspended in 50 μL RNase-free water and incubated for 10 minutes at 55°C. The concentrations of RNA were measured using a Smart-Spec plus spectrophotometer (Bio-Rad). Then, cDNA was synthesized from 0.5-1 g of total RNA in a 20 μL volume using a High-Capacity cDNA Reverse Transcription Kit (Cat#: 4368814, Applied Biosystem, Thermo Fisher Scientific). 5 minutes at 25 °C, 30 minutes at 42 °C, and 30 minutes at 95 °C were used in the RT-PCR procedure. 2 μL cDNA was combined with SYBR^TM^ Green PCR Master Mix (Thermo Fisher Scientific) and loaded into the CFX Connect Real-Time PCR Detection System as technical duplicates (BioRad). Primer sets for a panel of pro-inflammatory marker genes were employed. These included IFN-α, IFN-β, IFN-γ, TNF-α, iNOS, CD11b, IL-1, IL-6, and IL-8. Primer sequences are listed in **Table S7**. The raw data from real-time PCR raw data were analyzed as previously described [48,49]. After normalization to 18S rRNA, we assessed the difference in relative gene expression between the infection and control groups using CT values normalized to 18S RNA. To determine statistical significance, a *t*-test was performed.

## Author contributions

**Conceptualization**: Xusheng Wang, Ramkumar Mathur, He Huang

**Data Curation**: He Huang, Zachary Even, Xusheng Wang, Ramkumar Mathur, Mikhail Golovko, Svetlana Golovko

**Formal Analysis**: He Huang, Xusheng Wang, Zachary Even, Alyssa Erickson, Ling Li

**Funding Acquisition**: Xusheng Wang, Ramkumar Mathur

**Investigation**: He Huang, Zachary Even

**Methodology**: He Huang, Zachary Even, Xusheng Wang, Ramkumar Mathur, Mikhail Golovko, Svetlana Golovko

**Project Administration**: Xusheng Wang, Ramkumar Mathur

**Resources**: Xusheng Wang, Ramkumar Mathur, Mikhail Golovko, Svetlana Golovko

**Software**: Xusheng Wang, Alyssa Erickson, Ling Li

**Supervision**: Xusheng Wang, Ramkumar Mathur

**Validation**: Xusheng Wang, Ramkumar Mathur, He Huang, Zachary Even

**Visualization**: He Huang, Xusheng Wang, Alyssa Erickson, Zachary Even, Ling Li

**Writing – Original Draft Preparation**: He Huang, Xusheng Wang

**Writing – Review & Editing**: He Huang, Xusheng Wang, Ramkumar Mathur, Zachary Even, Ling Li

## Supporting information

**S1 Fig.** Our proteomics profiling data demonstrated that ITY protein expression in D2 mice is significantly elevated compared to that in B6 ones in both high dose and low dose infection.

**S2 Fig.** DEPs detected in B6 mice after *S*. Typhimurium infection compared with the control group. A. DEPs from high dose infection versus control group in B6 mice. B. DEPs from high dose versus low dose infection group in B6 mice.

**S1 Table** Proteins identified and quantified by LC/LC-MS/MS

**S2 Table** DEPs in spleen among controls and treatments (B6 and D2 controls, B6 and D2 treated by high dose and low dose of *Salmonella*)

**S3A Table** DEPs in spleen between B6 and D2 treated by high dose of *Salmonella*

**S3B Table** DEPs in spleen between B6 controls and high dose treatment of *Salmonella*

**S3C Table** DEPs in spleen between high dose and low dose treatment of *Salmonella* on B6 mice)

**S4 Table** Co-expression clusters detected by WGCNA

**S5 Table** Enriched pathways in each cluster

**S6 Table** Enriched pathways in each module

**S7 Table** Primer sequences for the pro-inflammatory cytokines in RT-qPCR analysis.

